# Novel NGS Pipeline for Virus Discovery from a Wide Spectrum of Hosts and Sample Types

**DOI:** 10.1101/2020.05.07.082107

**Authors:** Ilya Plyusnin, Ravi Kant, Anne J. Jääskeläinen, Tarja Sironen, Liisa Holm, Olli Vapalahti, Teemu Smura

## Abstract

The study of the microbiome data holds great potential for elucidating the biological and metabolic functioning of living organisms and their role in the environment. Metagenomic analyses have shown that humans, along with e.g. domestic animals, wildlife and arthropods, are colonized by an immense community of viruses. The current Coronavirus pandemic (COVID-19) heightens the need to rapidly detect previously unknown viruses in an unbiased way. The increasing availability of metagenomic data in this era of next-generation sequencing (NGS), along with increasingly affordable sequencing technologies, highlight the need for reliable and comprehensive methods to manage such data. In this article, we present a novel stand-alone pipeline called LAZYPIPE for identifying both previously known and novel viruses in host-associated or environmental samples and give examples of virus discovery based on it. LAZYPIPE is a Unix-based pipeline for automated assembling and taxonomic profiling of NGS libraries implemented as a collection of C++, Perl, and R scripts.

## 1 INTRODUCTION

Our ability to produce sequence data in the rapidly growing field of genomics has surpassed our ability to extract meaningful information from it. Analyzing viral data is particularly challenging given the considerable variability in viruses and the low coverage of viral diversity in current databases. It is estimated that a vast majority of virus taxa are yet to be described and classified (Geoghegan and Holmes, 2017). This challenge is further complicated by the high evolutionary rate of viruses leading to emergence of new virus lineages and the relative scarcity of viral genetic material in metagenomic samples (Rose *et al*., 2016). Next-generation sequencing (NGS) is a high-throughput, impartial technology with numerous attractive features compared to established diagnostic methods for virus detection (Mokili *et al*., 2012). NGS-based studies have improved our understanding of viral diversity (Cantalupo *et al*., 2011). There is considerable interest within virology to explore the use of metagenomics techniques, specifically in the detection of viruses that cannot be cultured (Smits *et al*., 2015; Graf *et al*., 2016). Metagenomics can also be used to diagnose patients with rare or unknown disease aetiologies that would otherwise require multiple targeted tests (Pallen, 2014) or emerging infections for which tests are yet to be developed.

In recent years, the role of the bacterial microbiome in health and disease has been acknowledged and studied extensively (Kataoka, 2016; Biedermann and Rogler, 2015). Nonetheless, the influence of the viral constituent of the microbiome (i.e., virome) has received considerably less attention. Recent research has indicated that both pathogenic and commensal viral species can modulate host immune responses and thereby either prevent or induce diseases (Lim *et al*., 2015; Neil and Cadwell, 2018). Additionally, recent research has revealed modifications in the virome that are related to diseases such as acquired immunodeficiency syndrome and inflammatory bowel disease (Norman *et al*., 2015). Accordingly, there is a need to identify novel viruses that may be established pathogens and to define wider links of the virome with health and disease. Beyond humans, the veterinary, wildlife, arthropod and environmental viromes have large implications in e.g. animal health, zoonotic emergence, and ecosystem research that require new tools to understand and study the virosphere.

Bioinformatics pipelines and algorithms designed for the analysis of NGS microbiome data can be separated into three groups. The first group includes pipelines for virome composition analysis. These pipelines mine the relative abundance and types of viruses present in a given sample. These pipelines include VirusSeeker (Zhao *et al*., 2017), Viral Informatics Resource for Metagenome Exploration (VIROME) (Wommack *et al*., 2012), viGEN (Bhuvaneshwar *et al*., 2018), the Viral MetaGenome Annotation Pipeline (VMGAP) (Lorenzi *et al*., 2011) and MetaVir (Roux *et al*., 2014). The second group includes pipelines that are designed for bacterial composition analysis, such as MG-RAST (Meyer *et al*., 2008). Pipelines in the third group, such as MetaPhlan2 (Truong *et al*., 2015), Kraken2 (Wood *et al*., 2019) and Centrifuge (Kim *et al*., 2016a), can perform composition analysis for all known taxa. There are also a number of tools, pipelines, and algorithms for virus discovery, including Genome Detective (Vilsker *et al*., 2019), VIP (Li *et al*., 2016), PathSeq (Kostic *et al*., 2011), SURPI (Ho and Tzanetakis, 2014), READSCAN (Naeem *et al*., 2013), VirusFinder (Wang *et al*., 2013) and MetaShot (Fosso *et al*., 2017). Most of these tools depend exclusively on nucleotide-level sequence alignments and can detect viruses with highly similar sequences to a known virus. These limitations make it impossible to detect extremely divergent novel viral sequences that do not share nucleotide similarity to any known or existing viral sequence (Wang *et al*., 2013; Takeuchi *et al*., 2014). For these divergent novel sequences, it is critical to utilize amino acid-based comparison. Although amino acid-based comparison is computationally more challenging and is used by very few published methods, this approach can facilitate the detection of novel viruses.

The availability of robust bioinformatics pipelines for virome detection and annotation from NGS data continues to be one of the critical steps in many research projects. Pipelines are needed to efficiently detect viral sequences present in a complex mixture of host, bacterial, and other microbial sequences. The discovery of viral sequences depends on sequence alignment with other viral sequences in databases, as, in contrast to bacteria where 16S RNA is present in all taxa, viruses lack ‘explanatory genes’ found in all taxa.

Lazypipe offers several advantages to the existing methods for taxonomic profiling of viral NGS data. Lazypipe outsources homology search to a separate server eliminating the need to install and update local sequence databases. This is helpful in both reducing the workload on the user and ensuring that all the latest viral sequences are covered by the homology search. Additionally, this feature can significantly reduce the threshold for employing Lazypipe by the less technically savvy researchers. Lazypipe uses SANSparallel (Somervuo and Holm, 2015) to search for amino acid homologs in the UniProtKB database. Searching for homologs in the protein space is expected to retrieve more distant viral homologs than searches with nucleotide sequences (Zhao *et al*., 2017). Furthermore, SANSparallel is approximately 100 times faster compared to the blastp search (Somervuo and Holm, 2015), which is the default search engine employed by nearly all other annotation pipelines that search the protein space. Unlike most taxonomic profilers, Lazypipe assembles and annotates viral contigs, thus reducing the workload on the downstream analysis. Lazypipe implements a flexible stepwise architecture that allows re-execution of individual steps or parts of the analysis. This architecture addresses the increased risk of execution failure that is inherent to the analysis of large NGS libraries. Lazypipe supports data formats that can be used both by human researchers and automated tools. Results are output in the form of intuitive excel tables and interactive graphs, but also, in the form of standardized taxonomic profiles that can be integrated with automated workflows.

Lazypipe does not perform direct taxonomic binning of reads, but instead links these to database sequences via the assembled contigs. This approach results in a very high accuracy of taxon retrieval (see Results), however, this may come at the cost of lower accuracy for read binning. The accuracy of read binning was not accessed in this work since the main objective was to construct a highly accurate taxonomic profiler. Still, we provide the option to retrieve reads linked to any reported taxon or contig. Lazypipe implements a simple and robust model for abundance estimation. In this model reads aligned to a given contig are equally distributed among the taxa found by the homology search. Although more elaborate models are certainly possible, our benchmarking suggest that this simple model is sufficient for adequate estimation of viral abundancies.

## 1 MATERIAL AND METHODS

### 1.1 Laboratory procedures and samples

The faecal sample from diarrheic American mink (*Neovison vison*) was collected in September 2015 from a fur production farm in Finland, as described previously (Smura *et al*.). The processing and sequencing of the sample were conducted using a protocol described in (Conceição-Neto *et al*., 2015).

The human patient samples were derived from the diagnostic unit of Helsinki University Hospital Laboratory (HUSLAB) and stored in −80° C. In this study, RNA was extracted using either QIAamp Viral RNA kit (Qiaqen Inc., Valencia, CA, USA) or EasyMag (bioMerieux) according to the manufacturer’s instructions, followed by real time PCR detection described in (Kuivanen *et al*., 2019) for TBEV, in (Haveri *et al*., 2020) for SARS-CoV-2, and in Smura et al (manuscript submitted) for entero- and parechoviruses. This study was done according to research permits HUS/32/2018 § 16 for project TYH2018322 and HUS/44/2019 § 13 for projects TYH2018322 and M1023TK001.

Prior to sequencing, samples were treated with DNase I (Thermo Fisher, Waltham, USA) and purified with Agencourt RNA Clean XP magnetic beads (Beckman Life sciences, Indianapolis, USA). Ribosomal RNA was removed using a NEBNext rRNA depletion kit (New England BioLabs, Ipswich, USA) according to the manufacturers protocol. The sequencing library was prepared using a NEBNext Ultra II RNA library prep kit (New England BioLabs).

Libraries were quantified using NEBNext Library Quant kit for Illumina (New England BioLabs). Pooled libraries were sequenced on an Illumina MiSeq platform, using either MiSeq v2 reagent kit with 150 base pair (bp) paired-end reads or a MiSeq v3 reagent kit with 300 bp paired-end reads.

### 1.2 Unix pipeline for assembly, taxonomic profiling and binning of NGS data

We implemented a UNIX pipeline for automated assembly and taxonomic profiling of NGS libraries. The pipeline also performs taxonomic binning of the assembled contigs. The workflow of our pipeline is illustrated in Fig 1. Our pipeline was implemented as a collection of Perl, C++, and R programs with a command-line user interface written in Perl. Our implementation allows for execution of the whole pipeline with a single command or performing each analysis step separately. This allows for great flexibility when working with large NGS libraries. Each pipeline step is described in more detail below.

**Figure 1.**
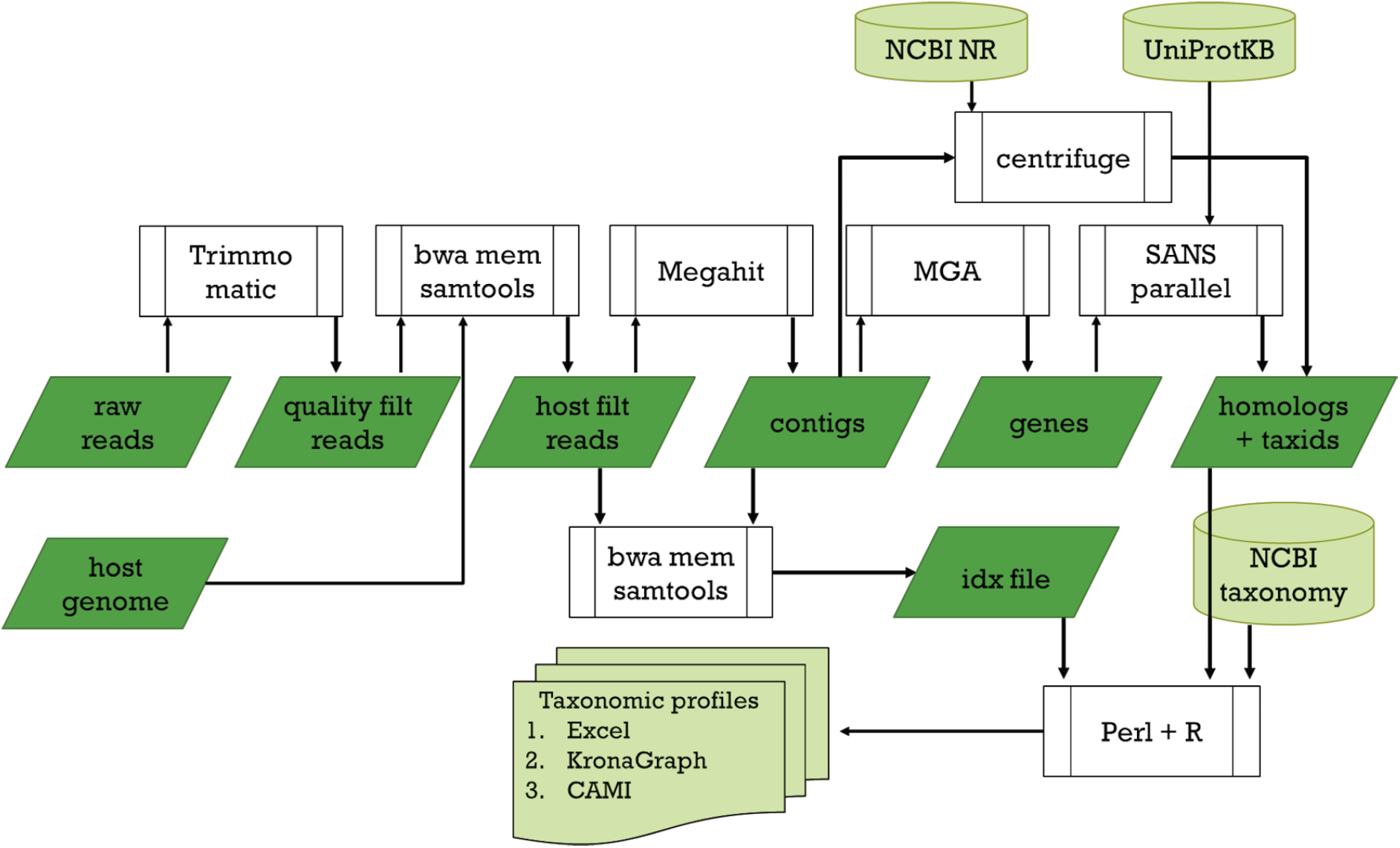
Lazypipe flowchart. Binaries and scripts are displayed in white, input and output files in green.

Paired-end libraries in FASTQ format (Cock *et al*., 2009) serve as input. First, primers, short reads and low-quality reads are removed with Trimmomatic (Bolger *et al*., 2014). Then host reads are filtered by aligning reads against the host genome with BWA-MEM (Li, 2013) and removing reads with high scoring alignments with SAMtools (Li *et al*., 2009). A threshold of 50 was selected by comparing the pre-assembly mapping of reads to the host genome to the post-assembly taxonomic binning of reads (without the host genome filtering). Comparing these for several samples showed that a threshold of 50 removes between 90% to 94% of reads that are assigned to *Eukaryota* in post-assembly binning while removing only 1% to 10% of reads that are assigned to Viruses (data omitted). Decreasing this threshold to 30 resulted in removal of only 63% to 64% of reads assigned to *Eukaryota* and increased removal of reads assigned to Viruses (18-29%). Increasing threshold to 100 again decreased the number of filtered eukaryotic reads (to 74-86%) with only slight improvement on the number of retained virus reads (0-9%). For the simulated metagenome (Fosso *et al*., 2017), setting the threshold to 50 results in filtering 99.93% of host reads and only 0.05% of viral reads (excluding the endogenous retroviral reads, which are filtered to a large extent). Thus, we selected 50 as a working threshold although we recognize that a more robust optimization can be performed.

In the next step reads are assembled with MEGAHIT (Li *et al*., 2015) or Velvet (Zerbino and Birney, 2008). MEGAHIT is used by default as this was the overall best assembler in the CAMI competition (Sczyrba *et al*., 2017). The pipeline then scans for gene-like regions in the assembled contigs with MetaGeneAnnotator (Noguchi *et al*., 2008) (default) or MetaGeneMark (Zhu *et al*., 2010) and translates these to amino acid sequences using BioPerl (Stajich *et al*., 2002). Extracted amino acid sequences are queried against UniProtKB using the SANSparallel (Somervuo and Holm, 2015) server. Top hits that pass a bitscore threshold value are used to assign contigs to the NCBI taxonomy ids. Note that contigs with several genes can be assigned to several taxonomy ids. We also support an alternative strategy of mapping contigs directly against the NCBI nr database. This is done by querying contigs with Centrifuge against NCBI nr and using alignments that pass a threshold value for the alignment score to assign contigs to taxonomy ids. We refer to this alternative version of our pipeline as the Lazypipe-nt.

Reads that passed host genome filtering are realigned to contigs using BWA-MEM top hits that pass a pre-set threshold on the alignment scores. Read distribution tables are generated using SAMtools (Li *et al*., 2009).

Next, taxonomy links generated by SANSparallel and read distribution tables are processed into an abundance table, which summarizes the number of contigs and reads binned to each taxon. Contigs that are mapped to two or more taxa contribute a corresponding fraction (such as 50%, 33%, 25% etc.) of mapped reads to the corresponding taxa. The raw abundance table is converted to an Excel file (using R) with several spreadsheets, each providing a different view of the acquired data. These views include the abundance of virus taxa (excluding bacteriophages), bacteriophages, bacteria, eukaryotes and, optionally, other high-level domains. For each of these groups, abundances are reported at three taxonomic levels (family, genus, and species). This arrangement allows for a rapid overview of NGS results and convenient “zooming in” on the taxa of interest. Taxonomic abundances are also presented as an interactive Krona graph (Ondov *et al*., 2011), which supports dynamic exploration of abundancies across different taxa. We also convert taxonomic abundances to CAMI Profiling Output Format (Sczyrba *et al*., 2017). By providing standardised output we support benchmarking of our pipeline by unbiased third-party evaluation initiatives such as CAMI (Sczyrba *et al*., 2017). Standardised output also aims to support simple and stable integration in automated workflows. To simplify accessibility, contigs for different taxa are sorted into a directory structure that follows the taxonomic hierarchy. A summary table is printed that lists all contigs for viruses, bacteriophages, bacteria and eukaryotes along with hits from SANSparallel or Centrifuge search.

In taxonomic profiling reads and contigs from the least abundant taxa have the highest risk of being misclassified. The organizers of the first CAMI competition addressed this problem by removing the last percentile of read distributions assigned by the compared taxonomic profilers (Sczyrba *et al*., 2017). We implement a similar strategy by assigning each taxon a cumulative frequency distribution value (*csum)*, which sums read frequencies mapped to that taxon and the more abundant taxa. We also assign confidence scores based on the csum score: the [0,95%] interval is assigned confidence 1, the [95%,99%] confidence 2 and the tail values [99%,100%] are assigned confidence 3. For a typical NGS library taxa with confidence score 1 will be true positives, those with score 3 (i.e. the last percentile) will be false positives and those with score 2 will represent borderline cases.

As an additional feature, Lazypipe offers an option to create interactive graphical reports that display the location and variation in viral contigs relative to reference viral genomes. This requires installation of a local database of viral reference genomes, which is then searched for taxa matching virus taxa found by the homology search. Contigs are aligned against the matching reference genomes with BWA-MEM and the resulting alignments are displayed with Integrative Genomics Viewer (Thorvaldsdóttir *et al*., 2013) in an internet browser.

In the last step we turn to quality control by generating graphical reports. The quality of the original library and assembly are monitored with histograms and key statistics. We also present the number of reads retained at consecutive pipeline steps: after quality filtering with Trimmomatic, after host genome filtering, after assembling and after gene detection. These are summarized as the survival-rate-plots.

### 1.3 Benchmarking performance

We evaluated our pipeline on the following two sets of data: a simulated metagenome from the MetaShot project (Fosso *et al*., 2017) and a mock-virome and bacterial mock-community data (SRA reference SRR3458569) (Conceição-Neto *et al*., 2015).

The MetaShot metagenome is a 20.5M PE 2×150 Illumina library simulated with ART [13]. We mapped reads in this library using accession numbers in read id-fields to 107 viral taxids, 99 prokaryote taxids and the human genome (94.5% of all reads). Strain taxids were further mapped using NCBI taxonomy to species, genus, family, order and superkingdom taxids resulting in 84 species and 46 genera of viruses, 71 species and 42 genera of bacteria. Based on this mapping we constructed a CAMI taxonomic profile (Sczyrba *et al*., 2017), which was then used as the gold standard in pipeline evaluation.

The mock-virome and bacterial mock-community is composed from 9 virus cultures (*Porcine circovirus* 2, *Feline panleukopenia virus, BK virus, Pepino Mosaic virus, Rotavirus A, Feline infectious peritonitis virus, Bovine herpesvirus 1, Dickeya solani LIMEstone bacteriophage and Acanthamoeba polyphaga mimivirus*) and 4 bacterial cultures (Conceição-Neto *et al*., 2015). The NGS library contains 12.4M PE Illumina HiSeq reads.

We compared the performance of Lazypipe on the MetaShot benchmark against Kraken2 (Wood and Salzberg, 2014), MetaPhlan2 (Truong *et al*., 2015), and Centrifuge (Kim *et al*., 2016b). Lazypipe was run with SANSparallel (referred to as *Lazypipe)* and Centrifuge (*Lazipipe-nt)* search engines. Kraken2, MetaPhlan2, and Centrifuge were run with default settings. For Centrifuge we used the NCBI nucleotide non-redundant sequences database; alignments with <60 nt match were removed to improve precision. Classification results were converted to CAMI taxonomic profiles and evaluated against the golden standard using OPAL: a CAMI (Sczyrba *et al*., 2017) spinoff project implementing CAMI metrics for metagenomic profilers (Meyer *et al*., 2019). For the simulated metagenome, we separately evaluated the entire taxonomic profile output by each of the pipelines and subprofiles limited to virus taxa.

## 2 RESULTS

### 2.1 Excellent recall and precision for both simulated and real datasets

Results for OPAL evaluation on the MetaShot benchmark are available from the project’s website (https://www.helsinki.fi/en/projects/lazypipe). Precision, recall (syn. sensitivity) and F1-score (harmonic mean of precision and recall) for predicted virus taxa and for all predictions are listed in Tables 1 and 2, respectively. For predicted virus taxa both Lazypipe variants have very high precision and recall at both genus and species level (Table 1). Lazypipe-nt has clearly the best balance between precision and recall, which is reflected in the highest F1-scores among the compared tools (Table 1). In the comparison of all predictions Lazypipe has the highest F1-score at the family and genus levels. Note that in this evaluation all methods have mediocre performance below the genus level. Lazypipe and Centrifuge are challenged with false positives and MetaPhlan2 and Kraken2 with false negatives (Table 2).

**Table 1.** Accessing accuracy of virus taxon retrieval by different tools.

| Tool | Rank | TP | FP | FN | Pr | Rc | F1 score |
| --- | --- | --- | --- | --- | --- | --- | --- |
| Lazypipe-nt | genus | 43 | 1 | 3 | 0.977 | 0.935 | 0.956 |
| Lazypipe |  | 41 | 1 | 5 | 0.976 | 0.891 | 0.932 |
| Centrifuge |  | 46 | 7 | 0 | 0.868 | 1.000 | 0.929 |
| MetaPhlan2 |  | 33 | 3 | 13 | 0.917 | 0.717 | 0.805 |
| Kraken2 |  | 21 | 1 | 25 | 0.955 | 0.457 | 0.618 |
| Lazypipe-nt | species | 73 | 1 | 11 | 0.986 | 0.869 | 0.924 |
| Lazypipe |  | 71 | 8 | 13 | 0.899 | 0.845 | 0.871 |
| Centrifuge |  | 84 | 43 | 0 | 0.661 | 1.000 | 0.796 |
| MetaPhlan2 |  | 37 | 8 | 47 | 0.822 | 0.440 | 0.574 |
| Kraken2 |  | 16 | 1 | 68 | 0.941 | 0.190 | 0.317 |

Compared tools are ordered by the descending F1-score for virus taxa predicted for simulated metagenome (Fosso *et al.*, 2017). Abbreviations: TP, true positives, FP, false positives, FN, false negatives, Pr, precision, Rc, recall, F1-score, harmonic mean of precision and recall.

**Table 2.** Accessing accuracy of viral and bacterial taxon retrieval by different tools.

| Tool | Rank | TP | FP | FN | Pr | Rc | F1 score |
| --- | --- | --- | --- | --- | --- | --- | --- |
| Lazypipe | genus | 84 | 17 | 5 | 0.832 | 0.944 | 0.884 |
| MetaPhlan2 |  | 71 | 6 | 18 | 0.922 | 0.798 | 0.855 |
| Lazypipe-nt |  | 68 | 19 | 21 | 0.782 | 0.764 | 0.773 |
| Kraken2 |  | 50 | 3 | 39 | 0.943 | 0.562 | 0.704 |
| Centrifuge |  | 83 | 161 | 6 | 0.340 | 0.933 | 0.498 |
| MetaPhlan2 | species | 105 | 10 | 51 | 0.913 | 0.673 | 0.775 |
| Lazypipe |  | 143 | 111 | 13 | 0.563 | 0.917 | 0.698 |
| Lazypipe-nt |  | 107 | 64 | 49 | 0.626 | 0.686 | 0.654 |
| Kraken2 |  | 53 | 21 | 103 | 0.716 | 0.340 | 0.461 |
| Centrifuge |  | 131 | 466 | 25 | 0.219 | 0.840 | 0.348 |

Compared tools are ordered by the descending F1-score for all predictions for simulated metagenome (Fosso *et al.*, 2017). Abbreviations: TP, true positives, FP, false positives, FN, false negatives, Pr, precision, Rc, recall, F1-score, harmonic mean of precision and recall.

To evaluate the performance of Lazypipe on real data, we ran the Lazypipe analysis with default settings on the mock-community data (for results please see project’s webpage). Recovery of the 9 mock-community viral taxa was evaluated by manual inspection of Lazypipe summary tables. Lazypipe recovered all 7 eukaryotic viruses included in the mock-virome. Moreover, the correct eukaryotic viruses were the only eukaryotic viruses predicted for this data with acceptable confidence scores (scores 1 and 2; excluding score 3, which has a high risk of being false positive). Thus, we had 100% sensitivity and 100% precision for the eukaryotic viruses at the species level. Lazypipe also reported the *Dickeya LIMEstone* virus, but did not report the *Acanthamoeba polyphaga mimivirus*.

### 2.2 Benchmarking time efficiency

We compared the execution time of Lazypipe, Kraken2, MetaPhlan2 and Centrifuge on the MetaShot simulated metagenome on a GNU/Linux machine with 64 2300 MHz CPUs. All programs were run with 16 threads. The wall clock time in the order from the fastest to the slowest was: Kraken2 (2min 30sec), Centrifuge (21min 43sec), MetaPhlan2 (2h 21min 21sec) and Lazypipe (4h 31min 51sec). Comparing this order to Tables 1 and 2 we see a trade-off between accuracy and speed. The fastest tools (Kraken2 and Centrifuge) are the least accurate, and the most accurate tools (Lazypipe and MetaPhlan2) are the slowest. Although Lazypipe is about twice as slow as MetaPhlan2, it is more accurate and creates annotated assembly, which is not done by any of the compared tools. We also note that key subprograms employed by Lazypipe (i.e. BWA, Megahit, SANSparallel and SAMtools) have parallel implementation and are expected to have good scalability.

### 2.3 Novel virome sequences from mink faecal samples

In addition to the mock-community data we tested the performance of Lazypipe using real data from different sample types (cerebrospinal fluid, serum, faeces, tissue samples) derived from various host species.

Since the pipeline is designed also for the detection of unknown viruses, we explored various sources for virus discovery with a by default unknown viral diversity. As an example of searching for the causative agents of veterinary disease, we analyzed sequence data derived from a faecal sample of a mink with gastroenteritis manifesting as diarrhea. Altogether, Lazypipe detected multiple contigs that indicated the presence of virus genomes (see Table 3). Notably, one contig contained a large ORF that most likely represents a novel picorna-like virus (order *Picornavirales*) with only 30% amino acid identity to the closest match. In addition, partial genomes of a toti-like virus with 29-32% amino acid identity to Beihai toti-like virus 4 (contig length 3792) and with 38-48% aa identity to Hubei unio douglasiae virus 1 (contig length 3219) were detected together with smaller fragments of other yet unclassified viruses (see Table 3).

**Table 3.**
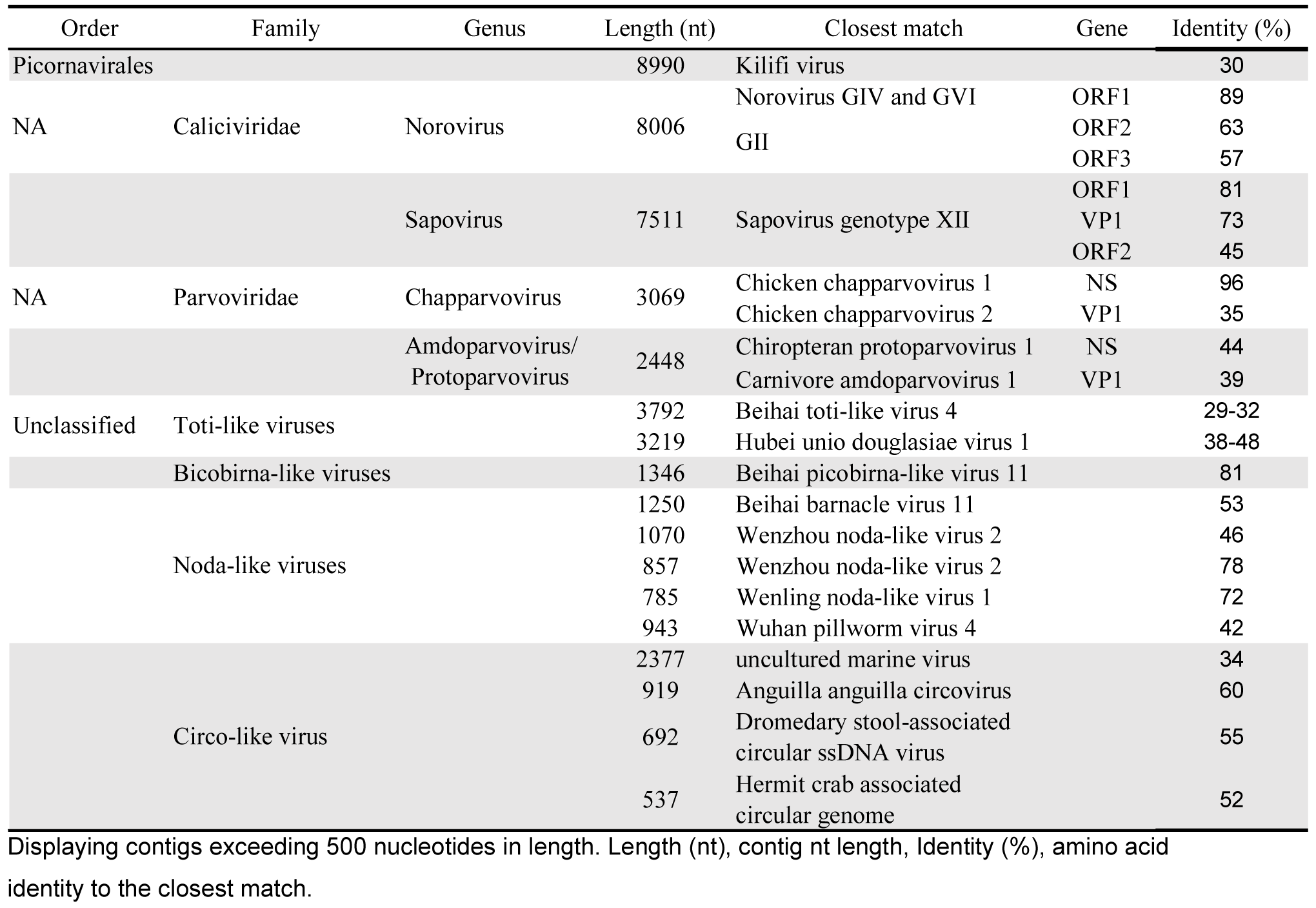
Virus contigs retrieved by Lazypipe for the mink fecal sample.

In addition to the above, virus groups with well-known association to the gastrointestinal system were detected. These included members of family *Caliciviridae* and *Parvoviridae*. Of the family *Caliciviridae*, six norovirus and six sapovirus contigs were detected. More thorough examination suggested that the norovirus contigs constitute a complete genome of a new representative of noroviruses with 89% amino acid identity in ORF1 (non-structural polyprotein) to norovirus genotypes IV and VI found in cats and dogs (Ford-Siltz *et al*., 2019), 63% amino acid identity in ORF2 (VP1) protein to genotype II found in pigs and 57% amino acid identity to genotype II in ORF3 (VP2).

The sapovirus contigs constituted a complete genome with 80-81% amino acid identity in ORF1 (including VP1 72-73% aa identity) and 45% amino acid identity in ORF2 (minor capsid protein VP2) to sapovirus GXII previously detected in minks (Oka *et al*., 2016; Guo *et al*., 2001). In addition to these, short low coverage contigs matching to Atlantic salmon calicivirus (78-100% amino acid identity) were detected. Most likely, these are derived from the feed.

Of the family *Parvoviridae*, the largest contig (3069 nucleotides) matched to chicken chapparvovirus 2 spanning from 3’end of the 5’end of VP1, whereas another large contig (2448 nucleotides) contained 3’end of NS protein with 44% aa similarity to Chiropterian protoparvovirus and the 5’ end of VP1 protein with 39% aa identity to Aleutian mink disease virus (amdoparvovirus). In addition to these, small fragments of mink bocaparvovirus were detected.

### 2.4 Human clinical samples

As an example of testing the suitability of Lazypipe for human clinical samples and exploring its use for detection of viral pathogens in humans, we used sequence data derived from human CSF, serum, brain tissue and nasopharyngeal swab samples that were previously tested positive for entero-, entero/parecho-, tick-borne encephalitis and SARS-coronavirus-2 viruses, respectively (Table 4).

**Table 4.**
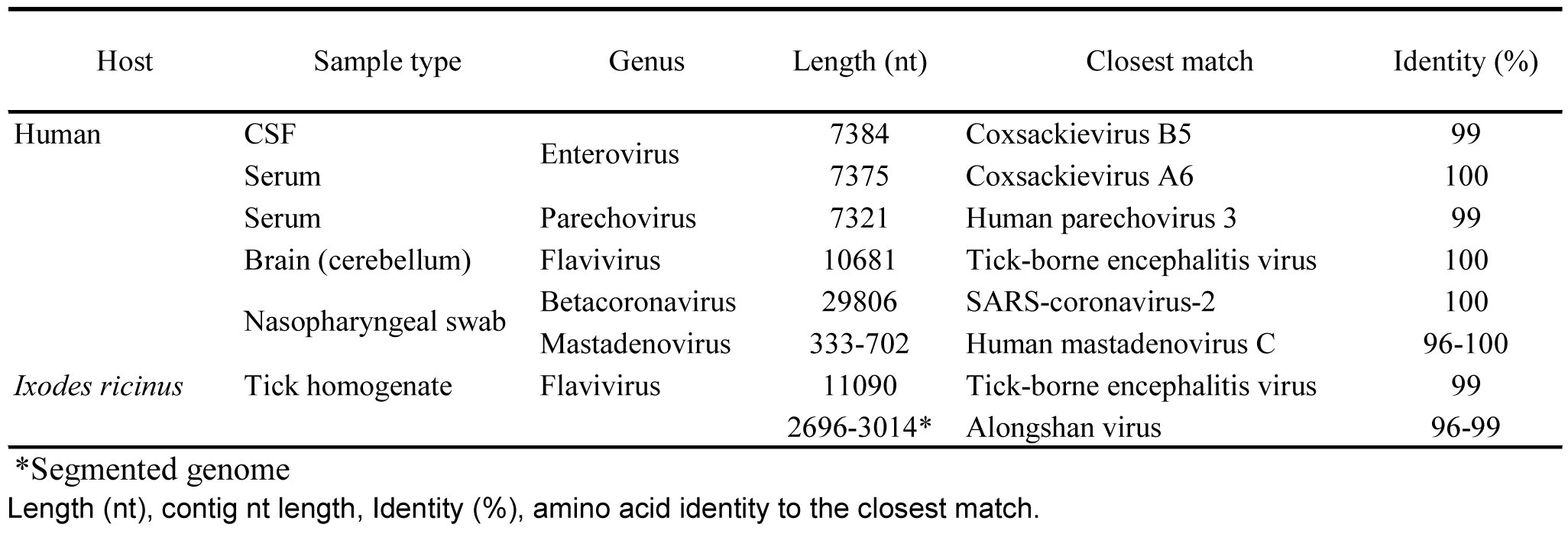
Lazypipe summary for various sample types with known human pathogenic viruses.

From a human CSF sample, a complete genome with 99% sequence identity with Coxsacievirus B5 (a member of Enterovirus B species) strains AU17EV1 and AU17EV2 (Queensland, Australia (Huang *et al*., 2017)) was retrieved. From two human serum samples complete genomes of Coxsackievirus A6 (Enterovirus A species) and Human Parechovirus 3 (Parechovirus A species) were retrieved. From a human cerebellum sample a complete genome of tick-borne encephalitis virus (TBEV) was retrieved. From a nasopharyngeal swab sample originated from the first case of COVID-19 in Finland (Haveri *et al*., 2020), a nearly complete SARS-coronavirus-2 (SARS-CoV2) genome and fragments of *Human mastadenovirus C* sequences were retrieved (see Table 4).

### 2.5 Arthropod samples

We also analysed samples of arthropod vectors. From an *Ixodes ricinus* tick sample collected from the Kotka archipelago in 2011, complete genomes of both Siberian subtype TBEV and a novel Alongshan virus (Kuivanen *et al*., 2019) were obtained (Table 4).

### 2.6 SARS-CoV2 patient samples from China

We analyzed public Illumina HiSeq/MiSeq libraries sequenced from bronchoalveolar lavage fluid from five patients with pneumonia at the early stage of the COVID-19 outbreak in Wuhan, China. Nine public NGS libraries were collected from NCBI SRA database (BioProject PRJNA605983) and analysed with Lazypipe. By applying default settings, we intentionally recreated a scenario, in which NGS data from SARS patients would be analysed prior to identifying the causative agent. *SARS-CoV* was identified by Lazypipe in all patients and in eight out of nine NGS libraries (Table 5). Lazypipe also identified co-infection with Influenza A in two out of five patients (Table 5).

**Table 5.** Lazypipe summary for SARS-CoV2 clinical samples from Wuhan, China.

| Accession | Library | Virus | Taxid | Readn | Readn% | Csumq | Contign |
| --- | --- | --- | --- | --- | --- | --- | --- |
| SRR11092058 | WIV02 | SARS-related coronavirus | 694009 | 36 | 0.517 | 3 | 9 |
| SRR11092063 | WIV02-2 | SARS-related coronavirus | 694009 | 559 | 0.368 | 1 | 23 |
| SRR11092057 | WIV04 | SARS-related coronavirus | 694009 | 732 | 13.088 | 1 | 15 |
| SRR11092062 | WIV04-2 | SARS-related coronavirus | 694009 | 5918 | 3.003 | 1 | 1 |
| SRR11092062 | WIV04-2 | Influenza A virus | 11320 | 274 | 0.139 | 1 | 2 |
| SRR11092062 | WIV04-2 | Autographa californica multiple nucleopolyhedrovirus | 307456 | 205 | 0.104 | 1 | 2 |
| SRR11092061 | WIV05 | SARS-related coronavirus | 694009 | 234 | 0.051 | 1 | 20 |
| SRR11092061 | WIV05 | Saccharomyces 20S RNA narnavirus | 186772 | 135 | 0.029 | 2 | 1 |
| SRR11092060 | WIV06-2 | SARS-related coronavirus | 694009 | 525 | 0.142 | 1 | 22 |
| SRR11092060 | WIV06-2 | Spodoptera frugiperda rhabdovirus | 1481139 | 165 | 0.045 | 1 | 1 |
| SRR11092060 | WIV06-2 | Saccharomyces 20S RNA narnavirus | 186772 | 103 | 0.028 | 2 | 3 |
| SRR11092064 | WIV07 | SARS-related coronavirus | 694009 | 56 | 0.070 | 1 | 9 |
| SRR11092059 | WIV07-2 | Influenza A virus | 11320 | 9063 | 0.097 | 1 | 4 |
| SRR11092059 | WIV07-2 | Saccharomyces 20S RNA narnavirus | 186772 | 3386 | 0.036 | 1 | 1 |
| SRR11092059 | WIV07-2 | SARS-related coronavirus | 694009 | 819 | 0.009 | 2 | 16 |
| SRR11092059 | WIV07-2 | Bamboo mosaic virus | 35286 | 325 | 0.003 | 2 | 1 |
| SRR11092059 | WIV07-2 | Spodoptera frugiperda rhabdovirus | 1481139 | 168 | 0.002 | 2 | 1 |

Lazypipe correctly identified SARS-CoV in 8 out of 9 samples. Additionally, two samples were identified with Influenza A and one sample with human mastadenovirus C coinfection. For details see the main text.

## 3 DISCUSSION

The availability of robust bioinformatics pipelines for viral metagenomics continues to be one of the critical steps in many research projects. Many of the existing pipelines are hindered by one or several limitations including large locally installed reference databases, slow homology search engines employed, low sensitivity for novel divergent sequences, low precision/recall performance for viral taxa or the lack of benchmarking for viral taxon retrieval, and the lack of assembling and contig annotation steps in the analysis. These limitations slow down the use of the unbiased sequencing approaches for rapid detection of novel emerging viruses.

In this publication we present Lazypipe, a novel bioinformatics pipeline that addresses the limitations typically encountered in viral metagenomics. Lazypipe avoids installation of large reference databases by delegating homology search to an external server. This frees the user from the need to install, index and update local reference databases, which can pose serious technical and resource constrains due to the sheer size of the modern sequence databases. By using SANSparallel (Somervuo and Holm, 2015) we also make Lazypipe considerably faster than pipelines based on BLASTP, and, simultaneously, render Lazypipe sensitive to highly divergent sequences, because viral peptides tend to be more conservative than the nucleotide sequences.

Taxonomic profiling by Lazypipe is done by querying assembled contigs instead of the reads, which translates into highly accurate taxonomic profiling of viral taxa. Benchmarking on simulated data showed that Lazypipe was clearly the most accurate taxonomic profiler for viral taxa among the four software packages compared. Testing on real mock community data demonstrated precision and recall nearing 100% for eukaryotic viruses. The detection of multiple novel viruses from various environmental and clinical samples reported here and in previous studies that used Lazypipe analysis (Kuivanen *et al*., 2019; Forbes *et al*., 2019) demonstrates that Lazypipe is also well suited for the detection and characterisation of novel and highly divergent viral genomes. Reflecting on the SARS-CoV2 pandemic situation (April 2020) we tested SARS-CoV2 positive Illumina libraries with Lazypipe and confirmed that the pipeline detected SARS-CoV in 9 out of 10 libraries with default settings and without SARS-CoV2 reference genome. This demonstrates the utility of Lazypipe for scenarios in which novel zoonotic viral agents emerge and can be quickly detected by NGS sequencing from clinical samples.

Previously, we have published two examples of novel and potentially zoonotic viral agents that were identified with Lazypipe from wild animals that can serve as vectors. A new ebolavirus was identified from faeces and organ samples of *Mops condylurus* bats in Kenya (Forbes *et al*., 2019), and a new tick-borne pathogen Alongshan virus from ticks in Northeast Europe (Kuivanen *et al*., 2019). These examples demonstrate the efficacy of Lazypipe data analysis for NGS libraries with very different DNA/RNA backgrounds, ranging from mammalian tissues to pooled and crushed arthropods.

The current pandemic highlights the need for an efficient and unbiased way to screen 1) for previously unknown viruses from either wildlife and arthropods for potential viral diversity that may emerge as human or animal pathogens, or 2) from individuals or human populations, production animals or companion animals manifesting with a disease of unknown aetiology for previously unknown or atypical causative agents. We showed here that Lazypipe can contribute to both of these important efforts and that it was able to detect the causative agent of the current pandemic without prior information.

## DATA AVAILABILITY

Lazypipe user manual and other resources are hosted at the project’s website (https://www.helsinki.fi/en/projects/lazypipe). Lazypipe source code is freely available from the Bitbucket repository (https://bitbucket.org/plyusnin/lazypipe/).

